# Accounting for uncertainty in residual variances improves calibration for fine-mapping with small sample sizes

**DOI:** 10.1101/2025.05.16.654543

**Authors:** William R.P. Denault, Peter Carbonetto, Ruixi Li, The Alzheimer’s Disease Functional Genomics Consortium, Gao Wang, Matthew Stephens

## Abstract

The Sum of Single Effects (SuSiE) model is a widely adopted method for genetic fine-mapping. We show that, in small-sample studies, the original SuSiE fitting procedure produces substantially higher rates of false positive findings. We show that a simple modification to SuSiE improves performance, especially in small-sample studies. This modification is particularly important for emerging molecular QTL applications in rare cell types and primary tissues where sample sizes are inherently limited.

## Main

Wang et al [1] introduced the “Sum of Single Effects” (SuSiE) model as a way to capture uncertainty in variable selection in multiple linear regression. This work was motivated by settings like genetic fine-mapping which often involves highly correlated variables, and where it is difficult to confidently identify the relevant variables, i.e., those with a non-zero regression coefficient. They pointed out that, even in such settings, it may be possible, and useful, to narrow down the potentially relevant variables by identifying sets of variables that, with high probability, contain at least one variable with a non-zero coefficient. They refer to these sets of variables as “credible sets” (CSs). The ability of SuSiE to efficiently generate accurate CSs is a key feature of the method. Since its introduction, a number of studies have extended the method in various ways [2–7].

Here we bring to light a limitation of the original SuSiE method: it can produce severely mis-calibrated CSs when applied to data sets with small sample sizes (*n* ≤50). We show that this miscalibration occurs even in the simplest case in which exactly one variable is assumed to have a non-zero coefficient. While many fine-mapping studies are large, this miscalibration issue is particularly important in smaller studies such as pilot experiments, complex experimental designs (e.g., dynamic eQTLs from [8], *n* = 19, and more recently [9], *n* = 53), costly multimodal-omics experiments (e.g., multimodal QTL mapping in influenza-infected cells [10], *n* = 35), and studies of rare cell types (e.g., eQTL mapping in microglia from the subventricular zone [11], *n* = 55).

To address this limitation, we show that the miscalibration can be improved by a simple modification of the procedures in [1] that accounts for uncertainty in the residual variance parameter. Specifically, whereas the original SuSiE estimates the residual variance by maximum-likelihood, here we instead average over the parameter under a diffuse prior, building on ideas from Bayesian statistics [12–15]. Briefly, the SuSiE model is based on a simpler “Single Effect Regression” (SER) model, which assumes that exactly one variable has a non-zero effect on the outcome. The original SER formulation does not account for uncertainty in the residual variance; here we extend the SER by placing a conjugate inverse-gamma prior on the residual variance. We refer to this extension as “SS-SER”, in which “SS” stands for “small sample”. The conjugate inverse-gamma setup here follows [13]. (We differ in using a proper Normal-Inverse-Gamma (NIG) prior with default 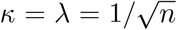 rather than the improper *κ*, →*λ* 0 limit they adopt, so that the posterior remains well-defined at small *n* while the prior’s influence shrinks with the sample size.) The SS-SER extension improves CS calibration in simulations, and generates more reasonable results in small-sample QTL studies, as we show here.

We compared our new approach (“SuSiE with SS-SER”) with the original approach (“SuSiE with default SER”) in simulated data sets. For each simulation, we sampled genotypes by randomly selecting loci from the 1000 Genomes Project, Phase 3 [16], including only unrelated individuals. Within these loci, we retained biallelic single nucleotide polymorphisms (SNPs) with minor allele frequencies (MAFs) ≥ 5%. From the remaining SNPs, we randomly selected 1–10 causal variants that collectively explained a specified proportion of the phenotypic variance (*h*^2^ = 10–75%). We also assessed the impact of different sample sizes (*n* = 20–200). For both methods, we set the upper limit on the number of causal SNPs, *L*, equal to the simulated number of causal variants, and used “purity filtering” [1] to filter out CSs containing weakly correlated SNPs (minimum absolute correlation of 0.5). More details are provided in Methods.

The results of these simulations show that, for small-sample studies (*n* ≤ 50), SuSiE with the default SER had consistently poor CS coverage, substantially below the target 95% level (Fig. 1, Supplementary Figures 1–5). By comparison, SuSiE with the SS-SER substantially improved coverage (although not attaining 95%). For example, with *n* = 20 and *h*^2^ = 25%, the default SER CS coverage was as low as 43%, whereas coverage did not drop below 59% using the SS-SER (Fig. 1). This improved coverage occured because the SS-SER produces larger CSs (Supplementary Fig. 3) with comparable purity (Fig. 1, Supplementary Fig. 4). Consequently, more CSs were discarded by the purity filter, resulting in a drop in power (Fig. 1, Supplementary Fig. 5). Although our assessments of fine-mapping performance have focussed on CSs because CSs account for uncertainty in the causal SNP due to LD, it is also common to assess performance of the posterior inclusion probabilities (PIPs), so for completeness we also assessed the PIPs produced by the two SuSiE variants (Supplementary Figures 6 and 7). The PIPs from the SS-SER variant of SuSiE always provided equal or better power at the same rate of false positives or false discoveries than the original SuSiE.

**Fig. 1.**
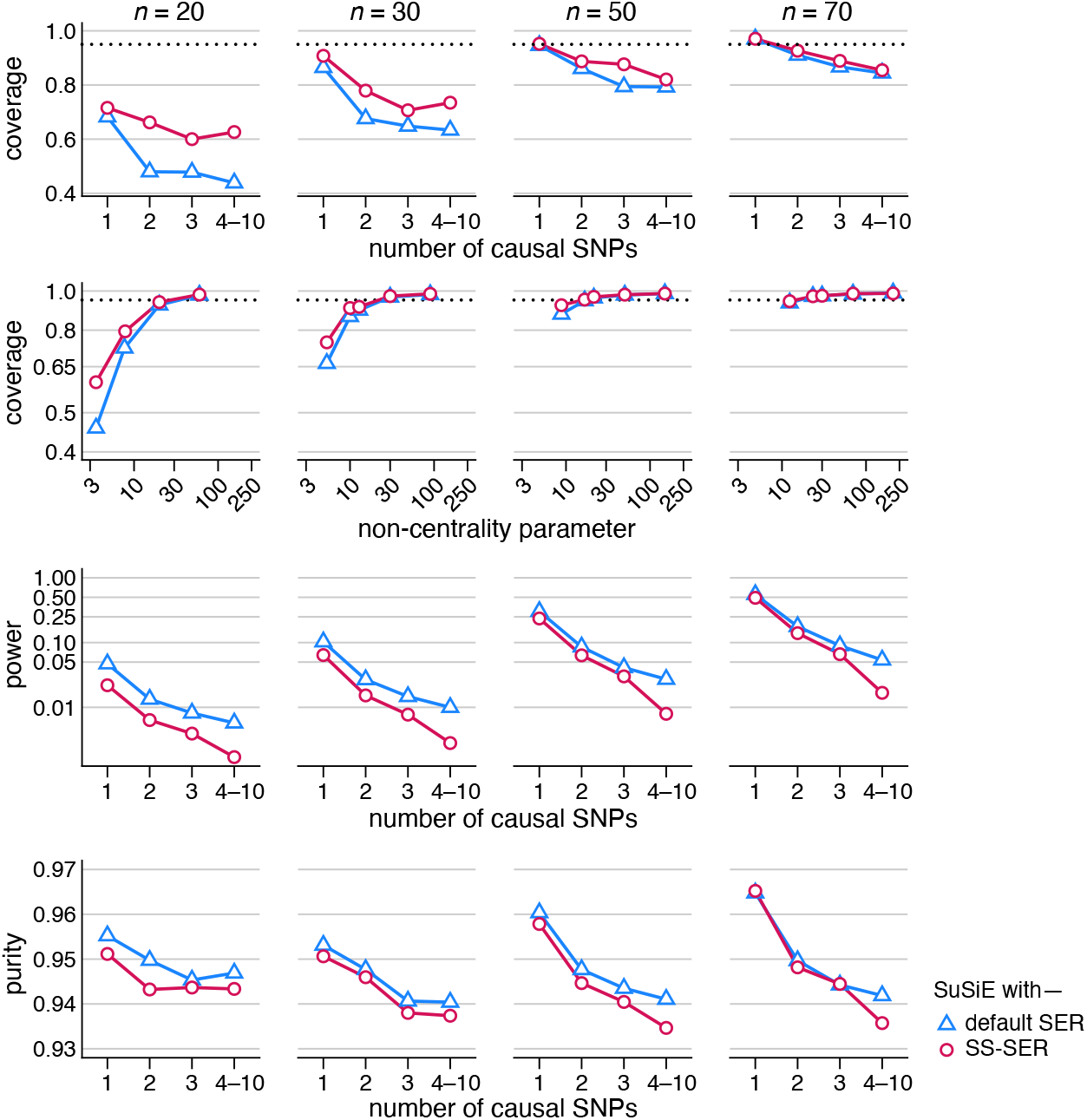
Comparison of credible sets (CSs) from SuSiE with the default SER and the “small sample” (SS) SER in simulations. In these simulations, genotype accounted for 25% of the trait variance (*h*^2^ = 25%). “Coverage” is the proportion of CSs that contain a true causal SNP; “purity” is defined as the smallest absolute correlation among all pairs of SNPs within a CS; and 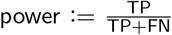, where TP, FN, TN denote, respectively, the number of true positives, false negatives and true negatives. “NCP” is the non-centrality parameter, NCP = *nh*^2^*/*(1–*h*^2^). The dotted black line represents the target coverage (95%). See also Supplementary Figures 1–5 for more results.

To disentagle the impact of estimating *L*—the number of causal SNPs or, equivalently, the number of CSs—on fine-mapping performance, we also conducted simpler simulations in which only a single variant affected the trait, and then we ran SuSiE by setting the upper bound on *L* to 1 (i.e., the true number) or to 10 (a considerable overestimate of the true number). We also ran SuSiE with and without filtering the CSs by purity. The results, summarized in the Supplementary Figures 8 and 9, show that overestimating *L* contributes to the miscalibration problem, but does not fully explain the extent of the miscalibration. This experiment also shows a broad improvement in calibration without the step of filtering CSs by purity, which would seem to suggest that one could improve calibration simply by omitting this step. However, without the purity filter, one typically ends up with many more “noninformative” CSs that trivially include many SNPs that are not the causal SNP (along with the causal SNP). Therefore, we generally recommend filtering CSs by purity.

To illustrate the impact of these ideas on real fine-mapping studies, we fine-mapped gene expression traits (eQTLs) from two RNA-seq experiments: one investigating transcriptional responses to primary macrophages before and after infection with influenza [10] (*n* = 35); and another studying eQTLs in microglia, a cell type important to brain aging [11] (*n* = 55). To finemap gene expression, we selected all SNPs with MAFs greater than 5% within ±200 kb of the gene’s transcribed region, and finemapped these SNPs, with *L*≤ 20. In the primary macrophages RNA-seq study, SuSiE with the default SER identified 772 causal SNPs (95% CSs) among the 13,680 that were analyzed, whereas SuSiE with the SS-SER more conservatively estimated 418 causal SNPs. Among the 394 genes in which SuSiE with the SS-SER identified exactly 1 causal SNP, the CS always overlapped a CS for the same gene that was identified by the original SuSiE. In the microglia RNA-seq study, SuSiE with the SS-SER was also generally much more conservative in identifying causal SNPs; for example, there were 3,523 genes identified by the original SuSiE as having 1 or more causal SNPs that had no causal SNPs under the SS-SER (Fig. 2A). Fig. 2B and C illustrate how “overfitting” to the small number of samples in this data set may contribute to poorly calibrated CSs, and shows that using the SS-SER remedies the issue somewhat.

**Fig. 2.**
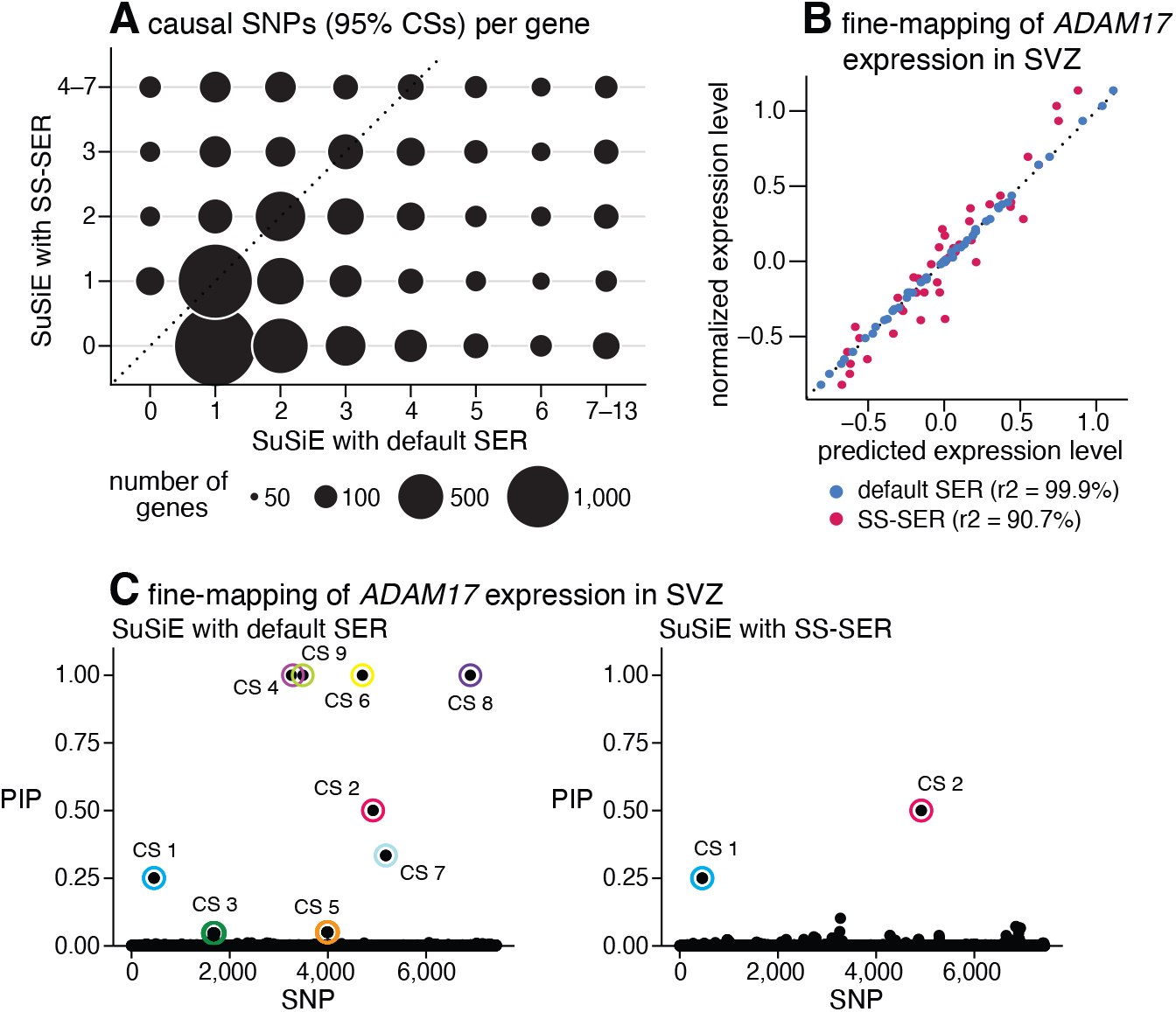
Fine-mapping of gene expression in the subventricular zone (SVZ) using RNA-seq data from microglial cells [11] (*n* = 55). **A**, number of causal SNPs (95% CSs) identified for each gene (one or both of the methods identified an expression SNP in 10,155 genes). **B**, SuSiE with the default SER appears to “overfit” a model of *ADAM17* expression (9 causal SNPs were identified), whereas SuSiE with the SS-SER also provides a strongly predictive model, but with only 2 causal SNPs. **C**, the posterior inclusion probabilities (PIP) and CSs for the *ADAM17* example; SuSiE with the default SER identifies 9 causal SNPs (CSs), whereas SuSiE with the SS-SER reports only 2 causal SNPs.

In conclusion, we have identified a limitation of SuSiE in which it produces poorly calibrated CSs in studies with small sample sizes, and we have developed a variant of SuSiE, “SuSiE with the SS-SER”, that improves calibration by modeling uncertainty in the residual variance parameter. Our method retains the computational advantages of SuSiE and can be applied to summary statistics [17], making it practical for a wide range of fine-mapping studies. While our approach improves CS coverage, some miscalibration remains in certain settings, and further improving performance for small-sample studies remains an area for future work.

## Methods

### Single effect regression accounting for uncertainty in residual variance estimates

The Sum of Single Effects (SuSiE) model is built on the simpler “Single Effect Regression” (SER) model, which assumes that *exactly one* of the variables in the regression has a non-zero effect on the outcome. The SER model from [1] is

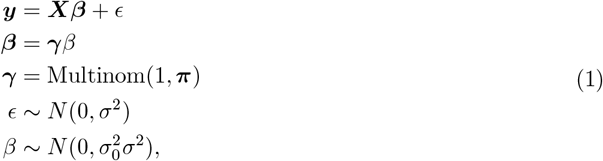

where ***y*** is the vector of observed outcomes, ***X*** is a matrix of explanatory variables, ***β*** is the vector containing the effects of these variables on the phenotype, and *ϵ* is Gaussian noise with variance *σ*^2^. Note that ***γ*** is a vector in which exactly one entry is non-zero. The prior probability for which entry is non-zero is encoded by the vector ***π*** (a vector of non-negative entries that sum to one). The hyperparameter 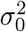 determines the prior on the effect size relative to the residual variance. In [1], the residual variance *σ*^2^ was estimated by maximizing the likelihood (or, in SuSiE, by maximizing the “evidence lower bound”). Given *σ*^2^ and 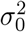, it is straightforward to compute the posterior distribution on ***β*** analytically, essentially by summing over all the possible values for ***γ*** [1, 13].

We note that (1) differs very slightly from the SER model in [1]; in [1], the prior variance on was 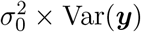, whereas here the prior variance is 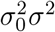. In most cases, we expect *σ*^2^ to be close to Var(***y***), so the practical effect of this change should be small.

To account for uncertainty in small-sample studies, we augment the SER model by placing a (conjugate) inverse-gamma prior on the residual variance,

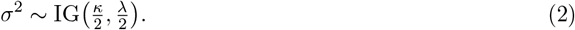

This results in a “Normal-Inverse-Gamma” (NIG) prior on *β, σ*^2^. We call this SER model the “SS-SER”, in which “SS” stands for “small sample”. The conjugate form follows [13, 14], who took the improper limit *κ, λ* → 0. We instead keep *κ, λ* strictly positive so that the marginal likelihood used to select 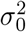 remains well-defined at small *n*, and set 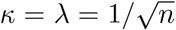 by default, so the prior is mildly informative at small *n* and fades into the data as *n* grows. Given *κ, λ* and 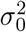, the posterior distribution on ***β*** can be computed analytically.

### Extension to the SuSiE model: SuSiE with the SS-SER model

The SER model is limited because it allows for only a single non-zero effect. Therefore, Wang et al [1] addressed this by allowing for *L*≥1 effects through summing *L* SER models, yielding the “sum of single effects” (SuSiE) model. They developed a fitting procedure, “iterative Bayesian stepwise selection” (IBSS), which fits the SuSiE model by iteratively fitting a sequence of SER models. They showed that this IBSS algorithm, when used with their SER, corresponds to optimizing a variational approximation [18] to the posterior distribution.

Here we use the same IBSS algorithm as in [1], but replace their SER (which fixes *σ* at a point estimate), with the SS-SER (which integrates out *σ*); see Supplementary Note for details. Unlike the original IBSS algorithm, this modified IBSS is not guaranteed to exactly optimize a variational approximation to the posterior distribution; however, our empirical results show that in small-sample studies it produces better CS calibration while maintaining the computational simplicity of the original IBSS algorithm.

### Univariate regression model

The derivations below hold for any *κ, λ >* 0, with the default 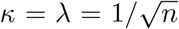 in our implementation; the same expressions remain valid in the improper limit *κ* = *λ* = 0.

Consider the following univariate regression model where *σ*^2^ is treated as unknown:

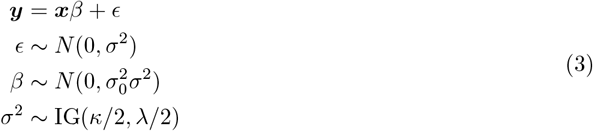

where ***y*** and ***x*** are in ∈ℝ^*n*^, *β* is an unknown parameter, *ϵ* is Gaussian noise, and 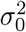 is a scaling factor on the effect size. IG(*κ, λ*) denotes the inverse gamma distribution with shape *κ* and scale *λ*.

The posterior distribution of *β* conditional on the other inputs and *σ*^2^ is

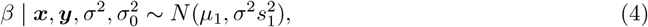

where

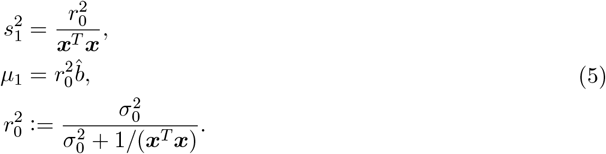

The posterior distribution of *σ*^2^ is

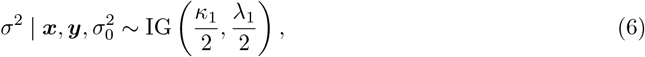

where

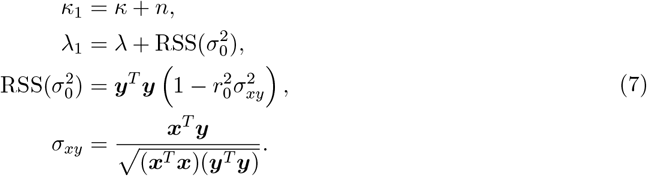

Thus, the posterior distribution of *β* is

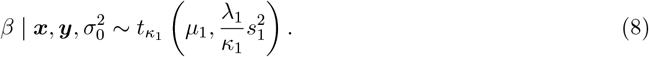

The Bayes factor (BF) for this model is given by

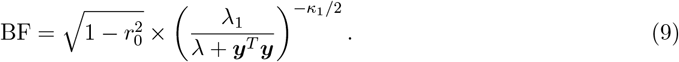

Further details of these derivations are provided in the Supporting derivations.

### Bayes factor and posterior distribution for SS-SER model

Under the SS-SER model (1–2), the posterior distribution of ***β***_*j*_ is a mixture of a point mass at zero and a *t*-distribution:

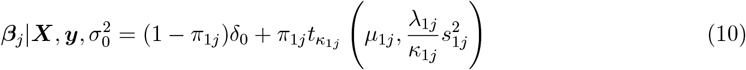

where 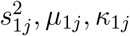, and *λ*_1*j*_ follow the same definitions as in the univariate case, substituting ***x*** with ***X***_*j*_ in equation (3). Here, *y* = ***X***_*j*_*β*_*j*_ + *ϵ*. The mixture weights are given by:

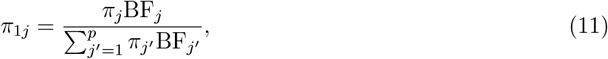

with where BF_*j*_ is the Bayes factor for the univariate association between column *j* of ***X*** and ***y*** (see eq. 9).

### Simulations

In all simulations, we randomly sample a genomic region from the 1000 Genomes Phase 3 dataset [16] and select *N* unrelated individuals within this locus. We retain only biallelic SNPs with a minor allele frequency (MAF) ≥5% among the selected individuals.

### Multiple causal SNPs simulations

For each simulation, we vary the number of individuals as *N* = 20, 30, 50, 75, and 100 and set the heritability, defined as the variance explained by the genotype, to *h*^2^ = 25%, 30%, 50%, and 75%.

For a given sample size *N* and heritability *h*^2^, we randomly select *N* individuals as described above. We then randomly determine the number of causal SNPs, *L*, where *L* is drawn uniformly from {1, …, 10}. The phenotype for individual *i* is generated as: *y*_*i*_ = *µ*_*i*_ + *ϵ*_*i*_, where 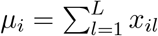. Where *x*_*il*_ denotes the genotype at the *l*^*th*^ causal SNP for individual *i*, and *ϵ*_*i*_ is a Gaussian noise term. To ensure that the in-sample variance explained by the genotype matches the specified *h*^2^, we rescale the error term *ϵ*_*i*_ to satisfy the following identity:

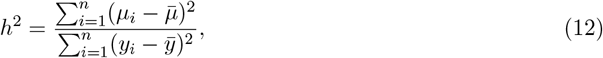

where 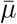 and 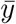 are the empirical means of *µ* and *y*, respectively. In these simulations, we set the upper limit on *L* to 10 when running SuSiE, regardless of the selected single-effect regression (SER) variant, and allow the method to estimate *L*.

#### Single causal SNP simulations

Simulations for the single causal SNP scenario follow the same setup as in the multiple causal SNP scenario, except that we fix the number of causal SNPs at *L* = 1. For each combination of *N* and *h*^2^, we evaluate the performance of SuSiE and its variants using the following configurations: (1) upper limit on *L* is 1, with or without purity filtering; (2) upper limit on *L* is set to 10, with purity filtering.

## Code availability

SuSiE with SS-SER is implemented in the original susieR package (https://github.com/stephenslab/susieR), demonstrated at https://stephenslab.github.io/susieR/articles/small_sample. The scripts to replicate these simulations are available at https://github.com/william-denault/susie_small_sample.

## Supplementary note

Here we provide detailed derivations of SuSiE with SS-SER model.

### Bayes factors

Consider the simple regression model to account for uncertainty in *σ*^2^

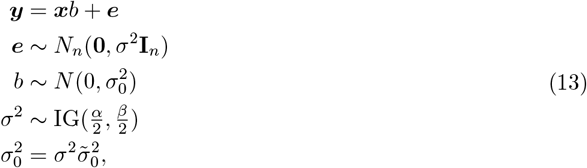

where IG(*a, b*) denotes the inverse gamma distribution with shape *a* and scale *b*, which has probability density function

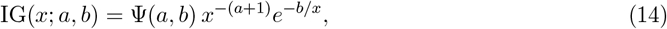

and we note 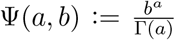 the *IG* normalizing factor. We note ***x*** := (*x*, …, *x*)^*T*^ ∈ **R**^*n*^, ***y*** := (*y*_1_, …, *y*_*n*_)^*T*^ ∈ **R**^*n*^ and ***e*** := (*e*_1_, …, *e*_*n*_)^*T*^ ∈ **R**^*n*^, *b* ∈ **R** is the regression coefficient, *σ*^2^ *>* 0 is the residual variance, 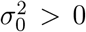 is the prior variance of the regression coefficient *b*, **I**_*n*_ denotes the *n* × *n* identity matrix, *N* (*µ, σ*^2^) denotes the univariate normal distribution with mean *µ*∈ **R** and variance *σ*^2^ *>* 0, and *N*_*d*_(***µ*, Σ**) denotes the multivariate normal distribution with mean ***µ*** ∈**R**^*d*^, and *d* ×*d* covariance matrix **Σ**. Here we assume that both ***x*** and ***y*** are centered to have mean zero, which removes the need for an intercept in the model [19]. In practice, we scale the prior variance of *b* by the sample variance of ***y*** to ensure that the prior is invariant to the scale of ***y***, but we can consider this to be “baked in” to the parameter 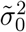, which is assumed to be known and is considered a fixed quantity in most of these derivations.

Given *σ*^2^ and 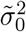, the posterior computations for this model are simple: they can be conveniently written in terms of the usual least-squares estimate of *b*, its variance, and the *z-*score, which are

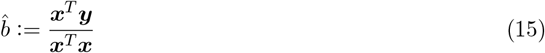

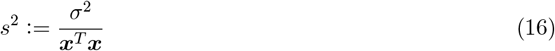

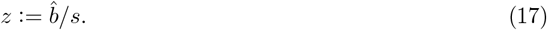

Note that scaling the prior variance of *b* by *σ*^2^ is important to obtain analytical expressions for marginal likelihoods and Bayes factors.

A few side notes:

a. Placing an inverse gamma prior on *σ*^2^ is equivalent to a gamma prior on the “precision” *σ*^2^ = 1*/σ*^2^ with shape *a* and inverse scale *b*.
b. The mean of IG(*a, b*) is 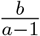 (for *a >* 1), its variance is 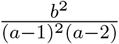 (for *a >* 2), and the mode is 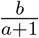.
c. The “noninformative” prior for *σ*^2^ is recovered with *α*→ 0, *β*→0, and this was the prior used in [14] and [13]. Instead, to keep the marginal likelihood used to select 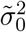 well-defined at small *n*, we take *α, β* strictly positive and use 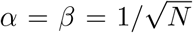 *N* as the default throughout this paper; the derived quantities remain valid in the improper limit *α* = *β* = 0.

From [13], the posterior distribution of *b* | *σ*^2^ is

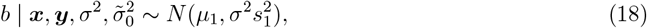

where

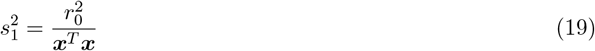

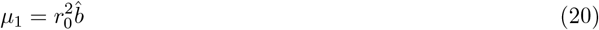

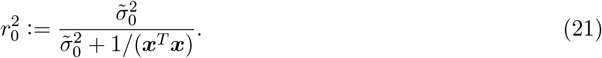

Writing the posterior mean and variance in this way is helpful for intuition: as *n* becomes large, the “prior” term 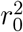 approaches 1 (because 1*/*(***x***^*T*^ ***x***) gets small), and therefore 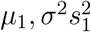 approach the least-squares estimate 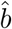 and its variance *s*^2^. Also note that when 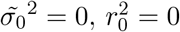.

The posterior distribution of *σ*^2^ | *b* is

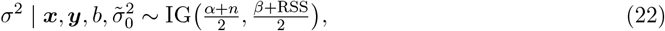

where 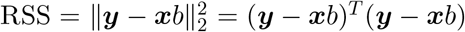 is the residual sum of squares. It is interesting to note that the posterior mode of *σ*^2^ when *α* = 0, *β* = 0 is simply RSS*/n*, the mean of the residual squared errors.

The posterior distribution of *σ*^2^ (averaging over *b*) is

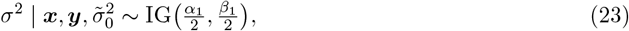

where

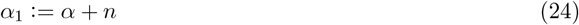

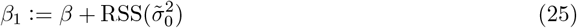

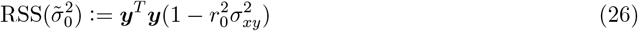

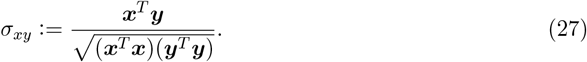

Here, 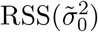 can be viewed as a Bayesian analog of the RSS [20]. This expression agrees with [13] (specifically, see eq. 4 in Protocol S1 of [13]). It is interesting to note that when *n* is large and *a* = 0, *b* = 0, the posterior mode of *σ*^2^ approaches 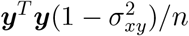, which is the sample variance of ***y*** scaled by a correction term for the correlation between ***x*** and ***y***. Also, when *α* = 0, *β* = 0, 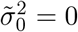, the posterior mean of *σ*^2^ is 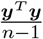.

Therefore, the posterior distribution of *b* is

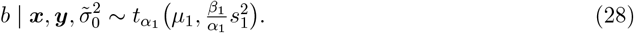

From this result, one can easily obtain the posterior mean and variance of *b*. Note that when *α* = 0, *β* = 0 and *n* is large, the posterior variance of *b* approaches

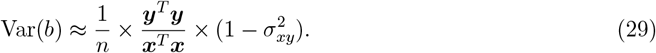

The Bayes factor for this model is

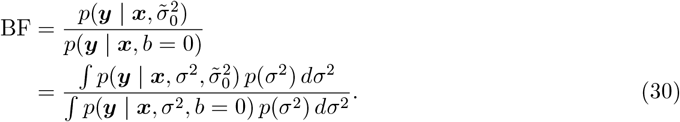

The dependencies on *α, β* are implicit in the expressions above and were not included to simplify notation. Let’s denote the marginal likelihood (integrated over *σ*^2^) as

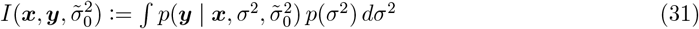

Then the Bayes factor for *b* in model 13 is

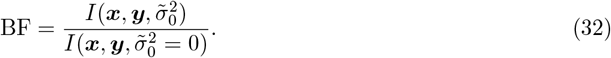

This integral works out to

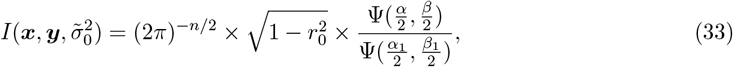

using the *α*_1_ and *β*_1_ defined in (24) and (25). When 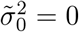, this simplifies to

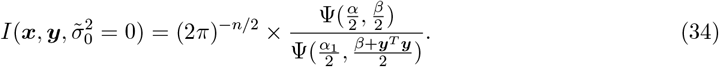

Therefore, the Bayes factor for this model is

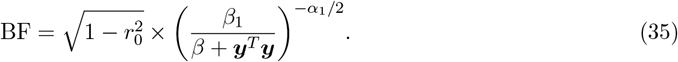

This expression agrees with the Bayes factor given in [13] (specifically, see eq. 12 in Protocol S1 of [13]). This also appears to agree with eq. 16 in [20].When *β* = 0, the Bayes factor simplifies to the somewhat intuitive expression

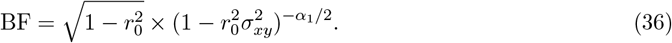

### Single effect regression

For convenience, we restate the single effect regression (SER) model from [1]. The SER extends (13) to a multiple linear regression model in which exactly one of the *p* variables affects the outcome, ***y***:

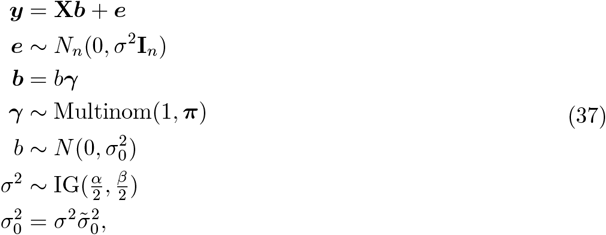

where **X** ∈ **R**^*n×p*^, ***y*** ∈ **R**^*n*^, ***γ*** ∈ {0, 1}^*p*^, and Mulitnom(*n*, ***π***) denotes the multinomial distribution with multinomial probabilities ***π*** = (*π*_1_, …, *π*_*p*_) and *n* trials. We assume ***y*** and the columns of **X** are centered to have mean zero, which implicitly accounts for an intercept. We will use ***x***_*j*_ to denote a column of **X**. Since our ultimate aim here is to extend this model to account for uncertainty, here we reparameterize the SER model slightly so that the prior variance is scaled by the residual variance, 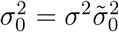. This has no effect on the derivations except for the part where we update *σ*^2^.

From our derivations above, it follows that the posterior distribution of *b*_*j*_, *σ*^2^ under the SER model has the following form [the dependence on *α, β* and ***π*** is implicit]:

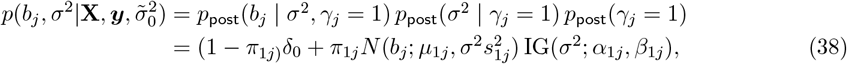

where 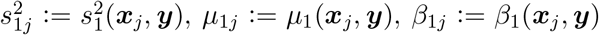, in which 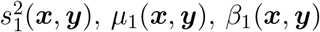 are given in equations (19, 20, 25). Here, we are reusing the expressions we have derived previously, with one change made to the notation to make explicit the dependence of these expressions on ***x, y***. As in the previous section, 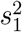 does not actually depend on ***y***, but we write it this way for consistency. The mixture weights are

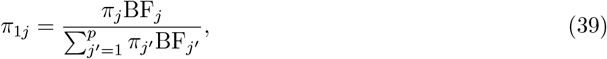

where BF_*j*_ := BF(***x***_*j*_, ***y***), and BF(***x, y***) is given by the previous result (35), again with the notation changed slightly to make the dependence on ***x, y*** explicit. And *α*_1*j*_ = *α* + *n* for all *j* = 1, …, *p*.

Following the previous notation we refer to SER as the function that outputs the quantities to compute the posterior quantities under the SER model with unknown *σ*^2^.

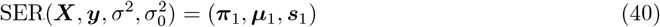

### Updating 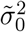

The only parameter that may need to be updated is the (unscaled) prior variance, 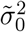.

The EM update for this parameter works out to

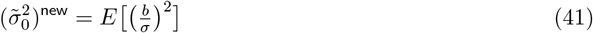

In other words, the EM update is given by the second moment of the coefficient, *b, scaled by the residual variance, σ*^2^. Expanding this, we have

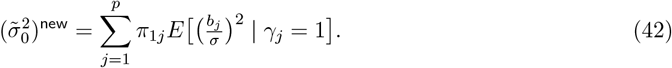

To derive the expected values in (42), let us refer back to the posterior distribution for the single regression model with unknown *σ*^2^. Let *t* = *b/σ*, and so we would like to find the mean and variance of *t*, because *E*[*t*^2^] = Var(*t*) + *E*[*t*]^2^. Recall, the posterior of *σ*^2^ was inverse gamma, *σ*^2^ ~ IG(*α*_1_*/*2, *β*_1_*/*2), and the posterior of *b* |*σ*^2^ was normal, 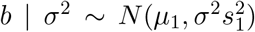. From the law of total expectation, and the fact that the posterior of *σ* is a *generalized gamma distribution* [21] (it is also a Nakagami distribution), the mean of *t* is

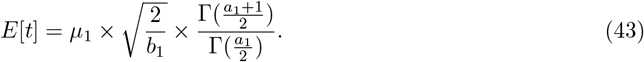

Also, using the law of total variance, the variance of *t* is

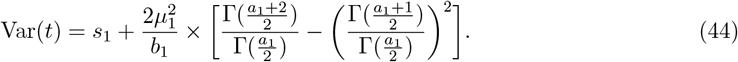

This completes the EM update for 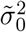.

Alternatively, 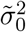 can be estimated by maximizing the marginal log-likelihood of the SER (which can be computed exactly) using a 1-d optimization algorirthm, e.g., using the `optimize()` function in R.

### Posterior for σ^2^

Following the fact that

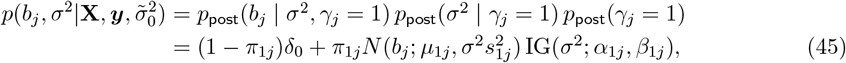

Assuming independence between on *p*_post_(*b*_*j*_ | *σ*^2^, *γ*_*j*_ = 1) leads to the following quantities

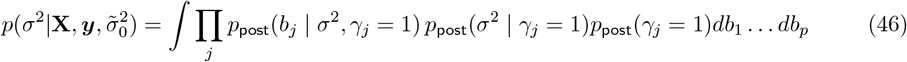

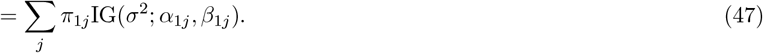

#### Algorithm 1

Generalized Iterative Bayesian Stepwise Selection (gIBSS) for the NIG prior

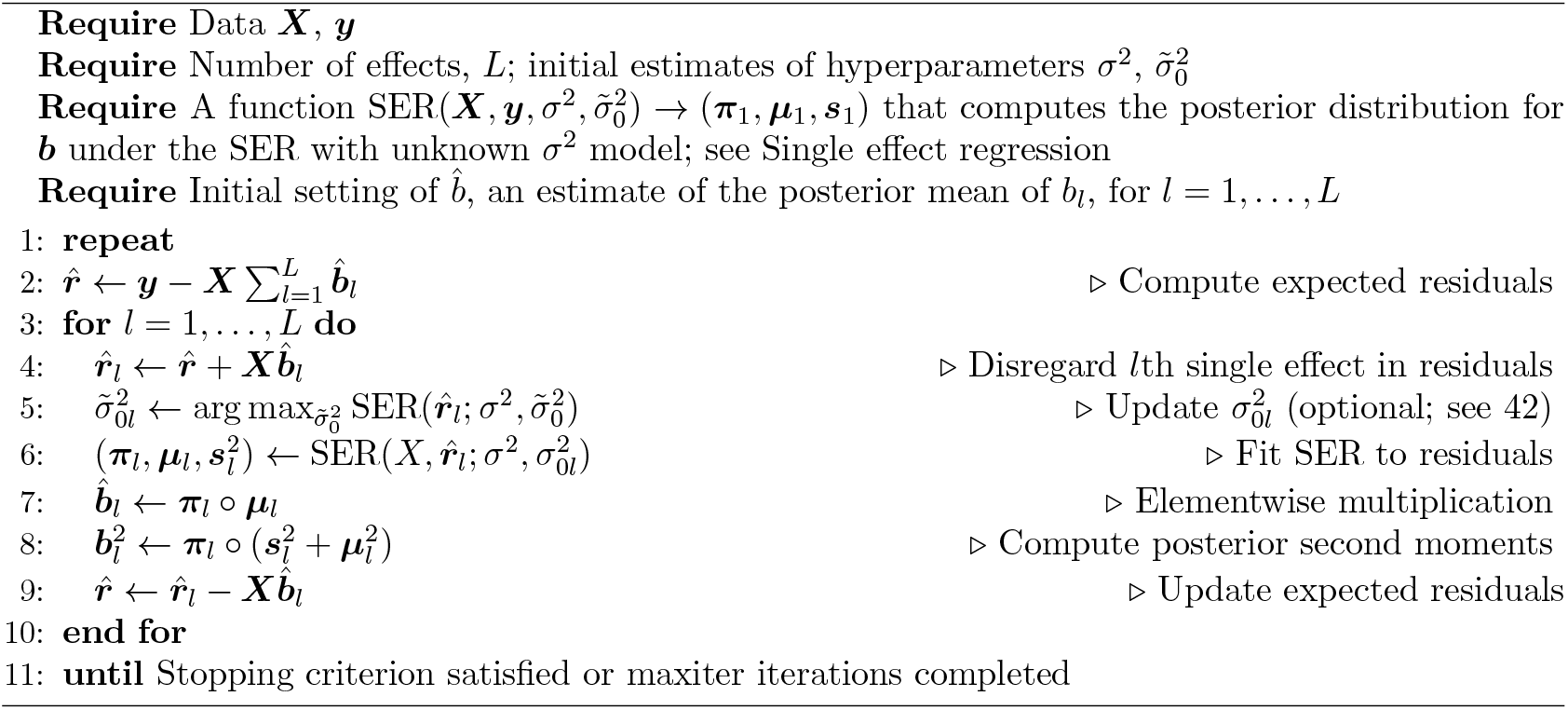

### The “Sum of Single Effects” (SuSiE) model with unknown σ^2^

We now extend the SER model with unknown *σ*^2^ to the “Sum of Single Effects” (SuSiE) model with unknown *σ*^2^. Instead of a single variable affecting the outcome ***y***, the SuSiE model allows at most *L* variables to affect ***y***:

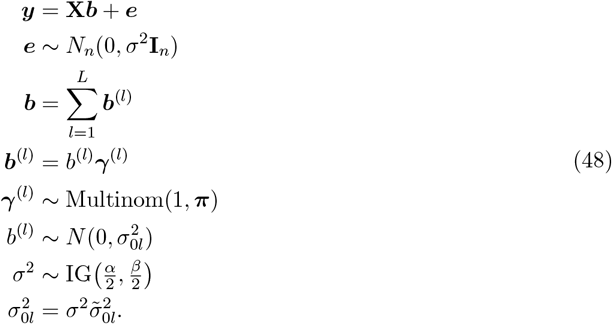

In the seminal SuSiE work [1], the authors assumed that the residual variance *σ*^2^ is either known or could be estimated via maximum marginal likelihood. They fit the SuSiE model using a variational approximation (VA) of the posterior, where the variational posterior assumes that the *L* single-effect coefficients are independent under the posterior:

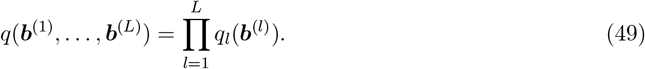

Wang and colleagues [1] further demonstrated that under this conditional independence assumption, each *q*_*l*_ must be a valid posterior distribution for the corresponding SER model. When extending SuSiE to the case where *σ*^2^ is unknown, the same conditional independence assumption leads to the following variational form:

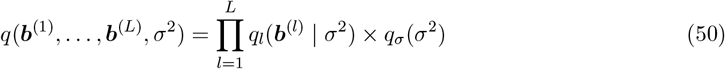

A key challenge in this variational approach is that *q*_*σ*_(*σ*^2^) turns out to be a mixture of *p* ×*L* inverse Gamma distributions, making it computationally intractable for efficient evaluation. However, given a known form for *q*_*σ*_(*σ*^2^), obtaining *q*_*l*_(***b***^(*l*)^ | *σ*^2^) is straightforward. Despite this, employing such a VA necessitates numerical shortcuts and additional approximations to maintain computational efficiency. To circumvent these computational difficulties, we adopt an alternative heuristic approach called Generalized Iterative Bayesian Stepwise Selection (gIBSS) [22] for fitting (48), as described below.

### gIBSS for SuSiE with unknown

*σ*^2^ A key feature of the (Gaussian SER) SuSiE model introduced by [1] is that, given estimates of *b*_1_, …, *b*_*L−*1_, estimating *b*_*L*_ reduces to fitting a single-SER model with an offset:

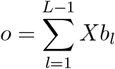

As noted by Wang and colleagues [1], this observation suggests an iterative approach to fitting the SuSiE model by repeatedly fitting SER models—an algorithm they refer to as Iterative Bayesian Stepwise Selection (IBSS). We build on this idea, augmenting SuSiE with the NIG prior using an iterative algorithm in which each iteration fits the SER with this prior. We refer to the resulting algorithm as Generalized IBSS (gIBSS), see Algorithm 1.

In short, gIBSS computes the approximate posterior distribution for each single effect *b*_*l*_, which allows computing 1) the posterior mean of each estimated effect and their corresponding credible set (CS). The CSs are computing following [1]. As opposed to the original IBSS, it is not proven that gIBSS optimizes a proper variational approximation of the posterior distribution of *b* under the NIG prior. Nonetheless, in our numerical experiment, gIBSS showed particularly good performances.

In summary, gIBSS computes an approximate posterior distribution for each single-effect coefficient *b*_*l*_, enabling the derivation of both (1) the posterior mean of each estimated effect and (2) the corresponding credible sets (CSs), which are constructed following [1]. Unlike the original IBSS, gIBSS is not proven to optimize a formal variational approximation of the posterior distribution of *b* under the NIG prior. Nevertheless, our numerical experiments demonstrate strong empirical performance of gIBSS.

### gIBSS stopping criterion

As described in Algorithm 1, gIBSS terminates when either the maximum number of iterations is reached (set to 100 by default) or a predefined stopping criterion is satisfied. In standard VA methods, it is common practice to monitor the evidence lower bound (ELBO) [18] and halt the algorithm when the increment in ELBO between consecutive iterations falls below a certain threshold (e.g., 10^−6^).

Since gIBSS does not explicitly optimize a VA objective, we do not have direct access to an ELBO. Instead, we assess the variation in the parameters (***π***_1_, …, ***π***_*L*_) across iterations and stop when the change between consecutive iterations is sufficiently small, i.e.,

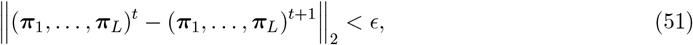

where *ϵ* is a small threshold, set to 10^−6^ by default. While this stopping criterion is heuristic, we note that some fine-mapping methods that explicitly optimize a variational approximation employ a similar approach in practice (see the implementation of SuSiE-inf [6]).

### Some notes on the t distribution

Let *x*∽ *t*_*ν*_ be random variable drawn from the *t* distribution with *ν* degrees of freedom. The probability density function is

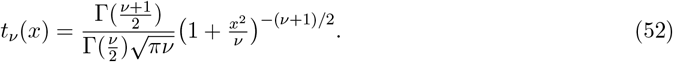

Now let *y* = *µ* + *σx*. We denote its distribution as *y* ~ *t*_*ν*_(*µ, σ*^2^), and its probability density function is

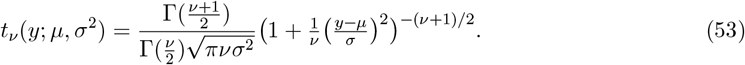

The mean of *y* is *µ* (for *ν* > 1) and its variance is 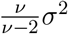(for *ν* > 2).

A useful identity: if 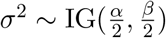 and 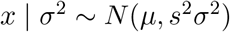, then 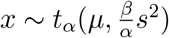.

### Posterior distribution of σ^2^, and b when σ^2^ is unknown

Let’s denote

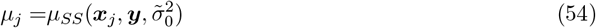

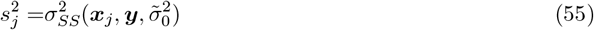

Then the posterior first and second moment of *b*_*j*_ under the SER SS model 37 is given by

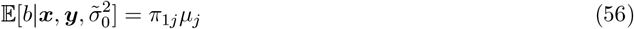

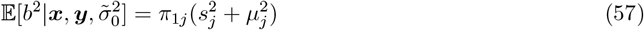

## Supplementary figures

**Supplementary Figure 1.**
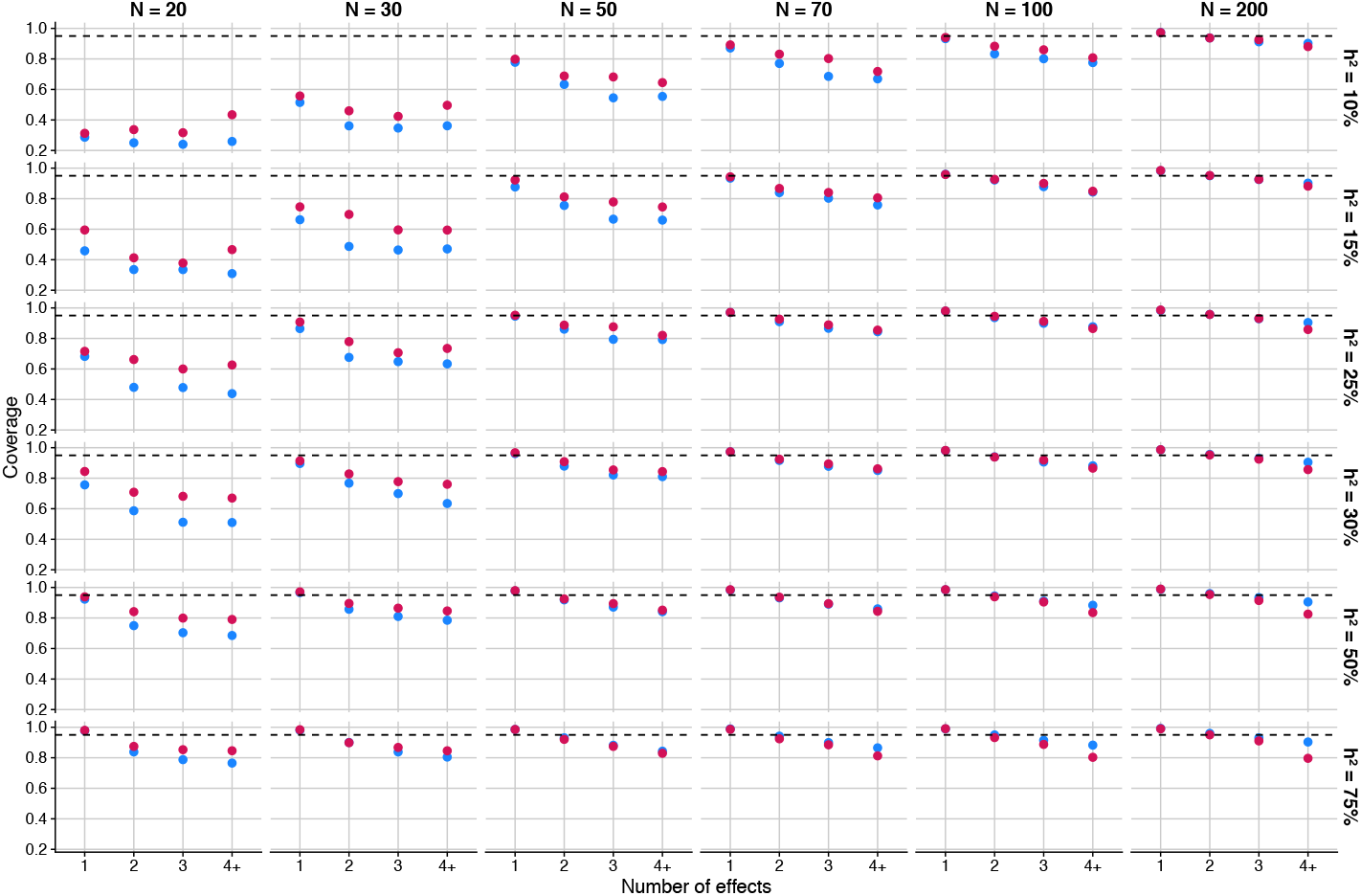
Credible set coverage in simulations. Blue = SuSiE with the default SER, red = SuSiE with the SS-SER. The dashed line gives the target CS coverage of 95%.

**Supplementary Figure 2.**
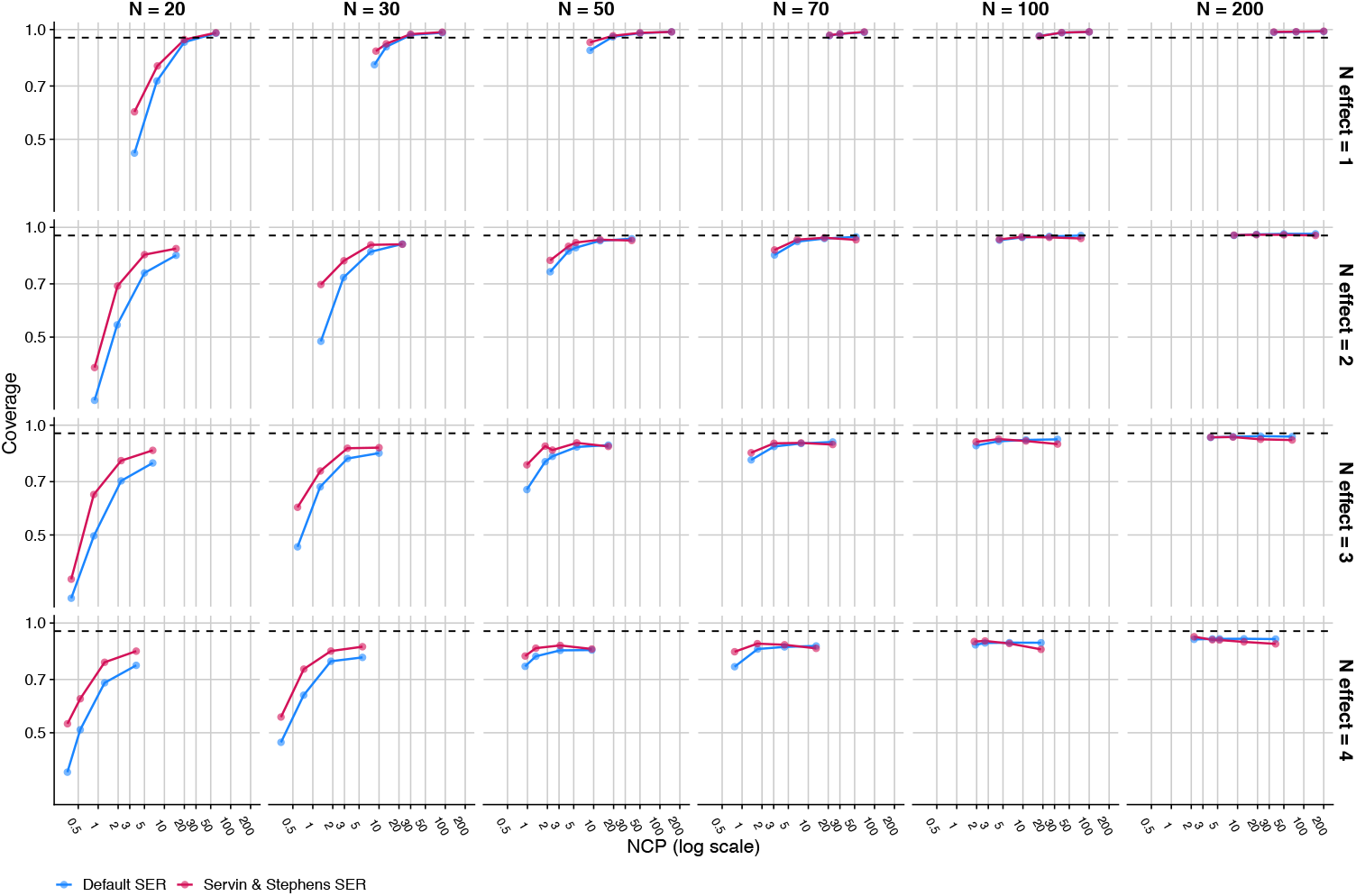
CS coverage vs. NCP in simulations. Blue = SuSiE with the default SER, red = SuSiE with the SS-SER. The dashed line gives the target CS coverage of 95%.

**Supplementary Figure 3.**
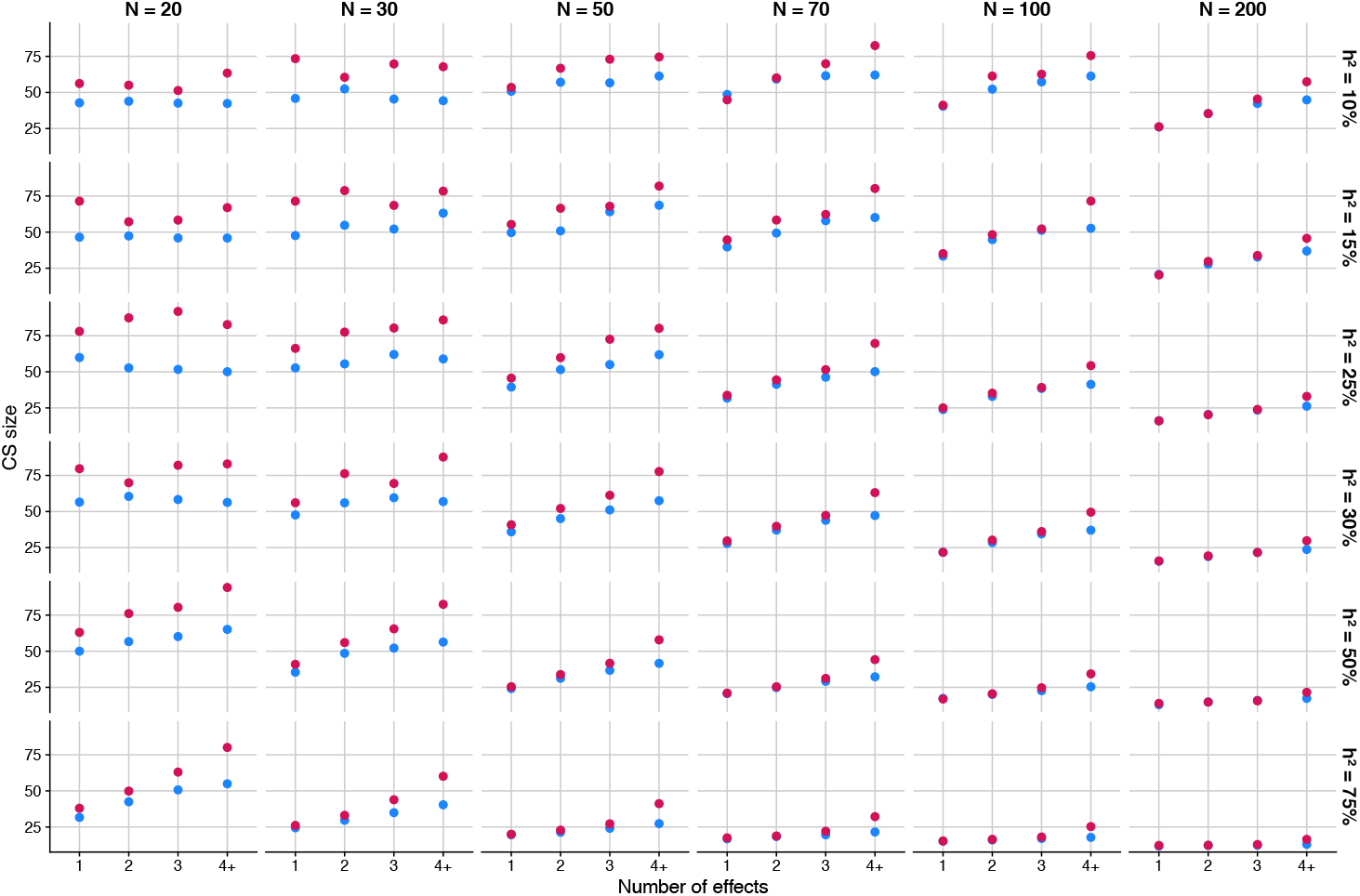
Mean credible set size in simulations. Blue = SuSiE with the default SER, red = SuSiE with the SS-SER.

**Supplementary Figure 4.**
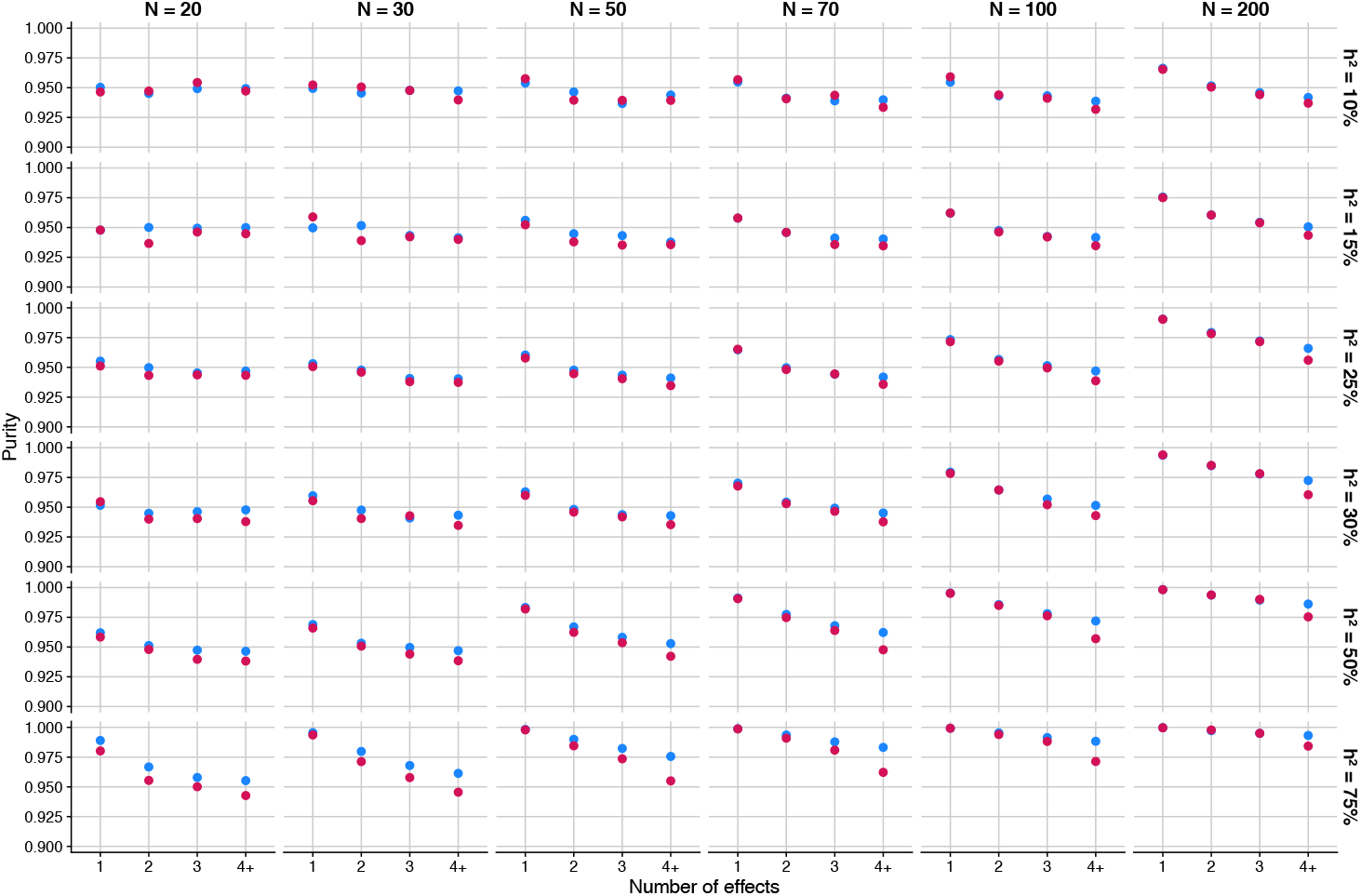
Credible set purity in simulations. Blue = SuSiE with the default SER, red = SuSiE with the SS-SER.

**Supplementary Figure 5.**
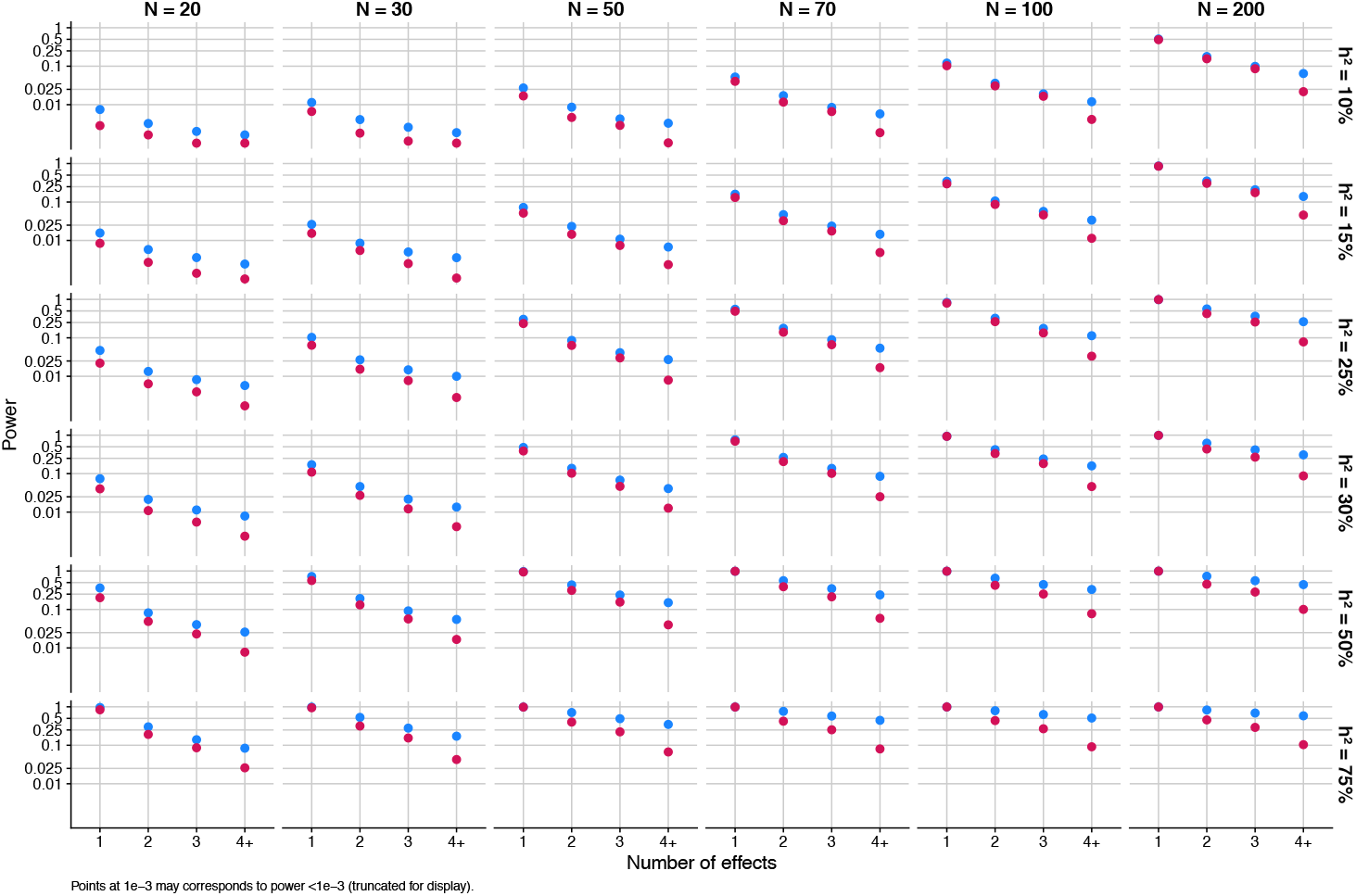
Credible set power in simulations. Blue = SuSiE with the default SER, red = SuSiE with the SS-SER. Note that power less than 0.001 is shown as 0.001.

**Supplementary Figure 6.**
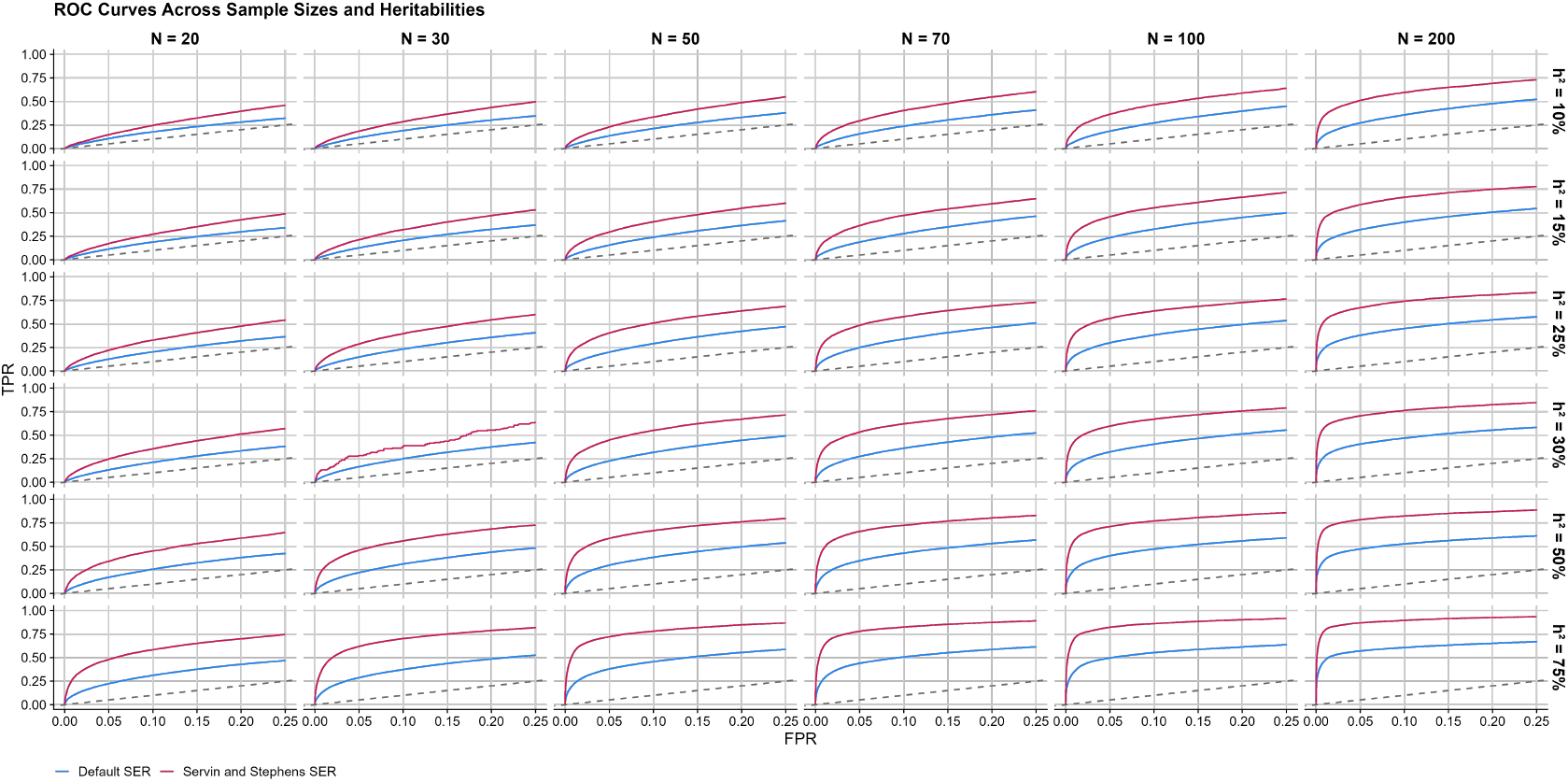
Detection of causal SNPs in simulations using PIPs: ROC curves. Blue = SuSiE with the default SER, red = SuSiE with the SS-SER. Note that the “true positive rate” (TPR) is the same as power.

**Supplementary Figure 7.**
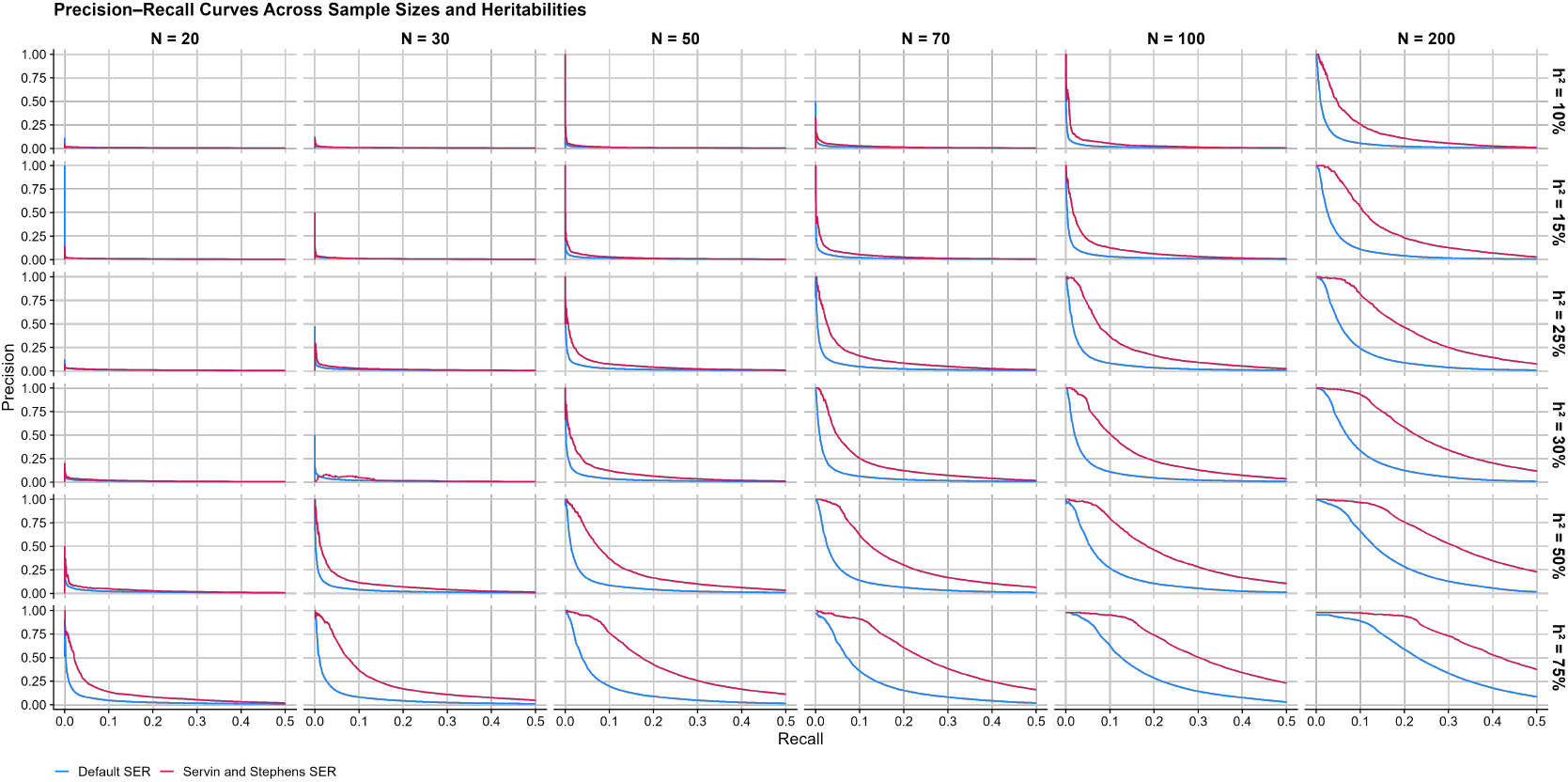
Detection of causal SNPs in simulations using PIPs: precision-recall curves. Blue = SuSiE with the default SER, red = SuSiE with the SS-SER. Note that these plots are the same as power-FDR curves after flipping the y-axis because 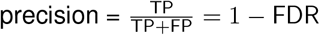 and recall = power.

**Supplementary Figure 8.**
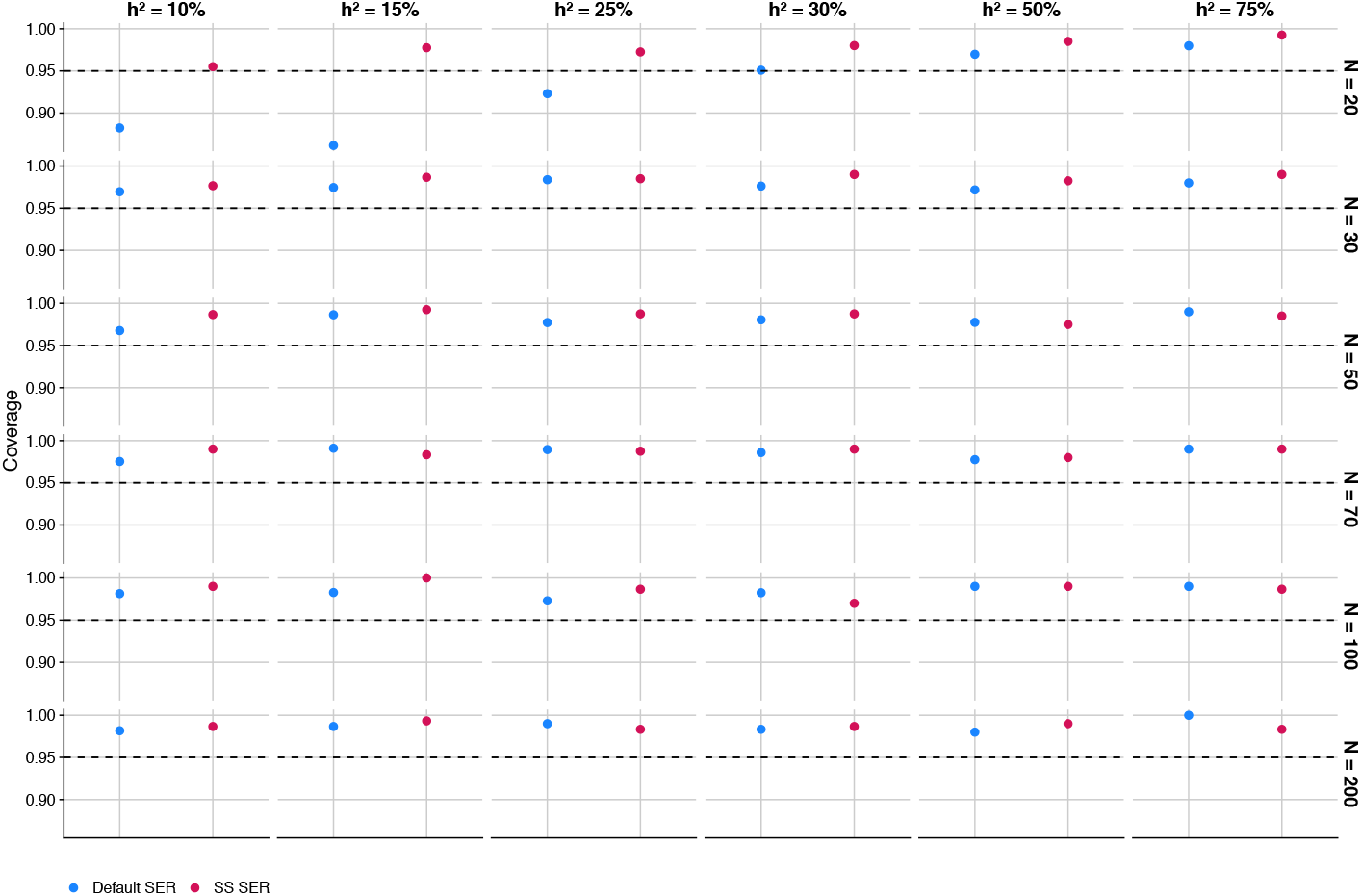
CS coverage in simulations with 1 causal SNP, without filtering of CSs by purity. The dashed line gives the target coverage level (95%).

**Supplementary Figure 9.**
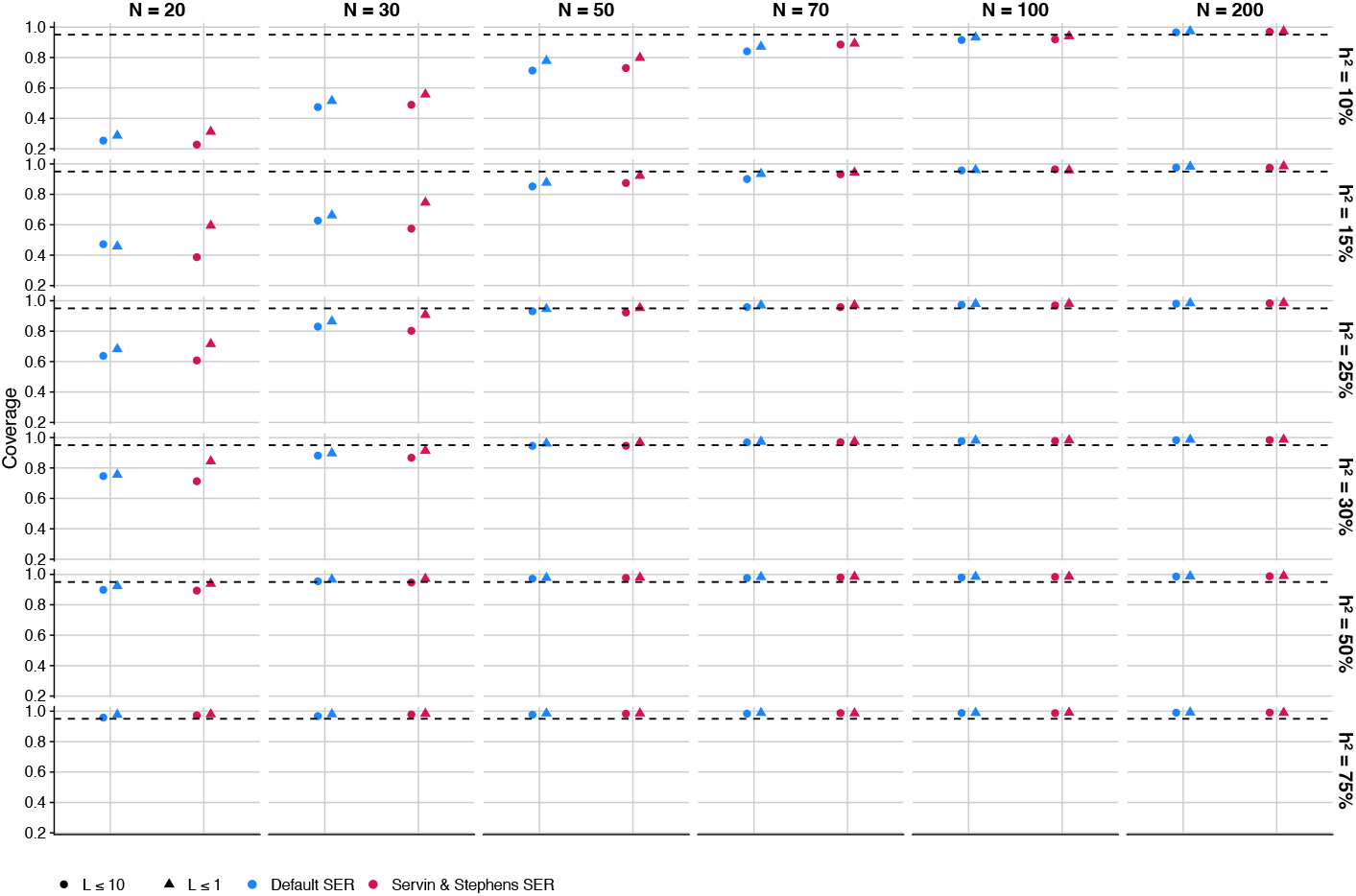
CS coverage in simulations with 1 causal SNP, with filtering of CSs by purity. The upper limit on the number of causal SNPs, *L*, was set to 1 or 10. The dashed line gives the target coverage level (95%).

